## Supplementary information for "*Vibrio cholerae* serotype impacts pathogenicity"

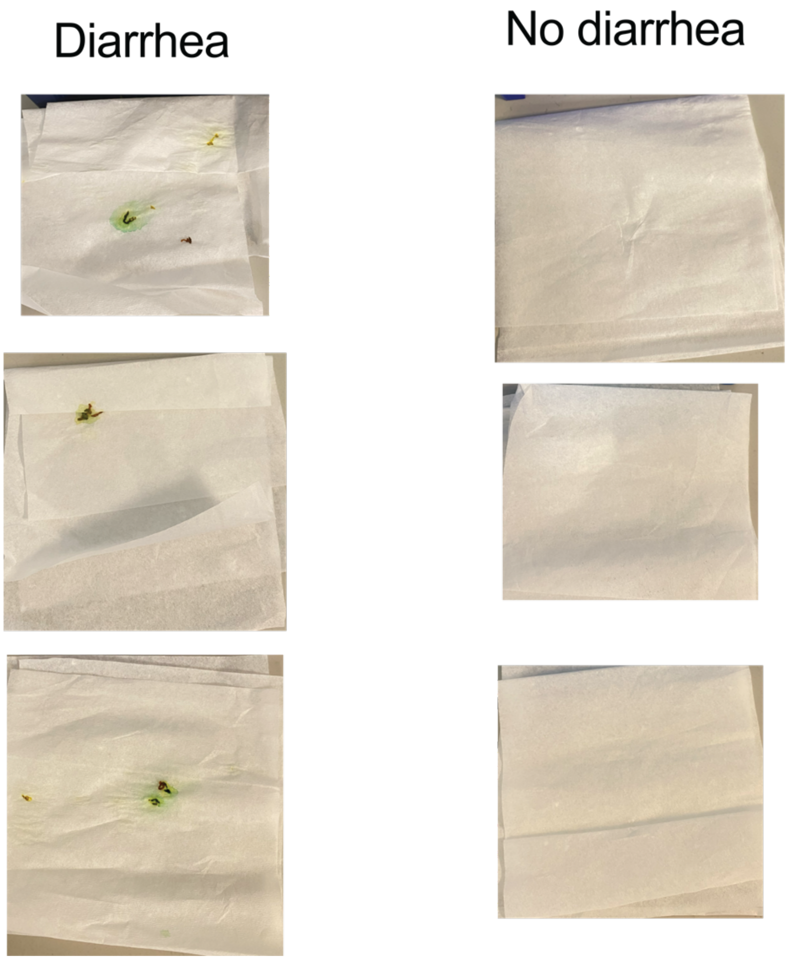


Supplementary Figure 1: **Indications of diarrhea.** Representative images of suckling mouse beddings identified with or without diarrhea. All mice used were Crl:CD1(ICR) mixed sex, postnatal day 5 at the time of infection.

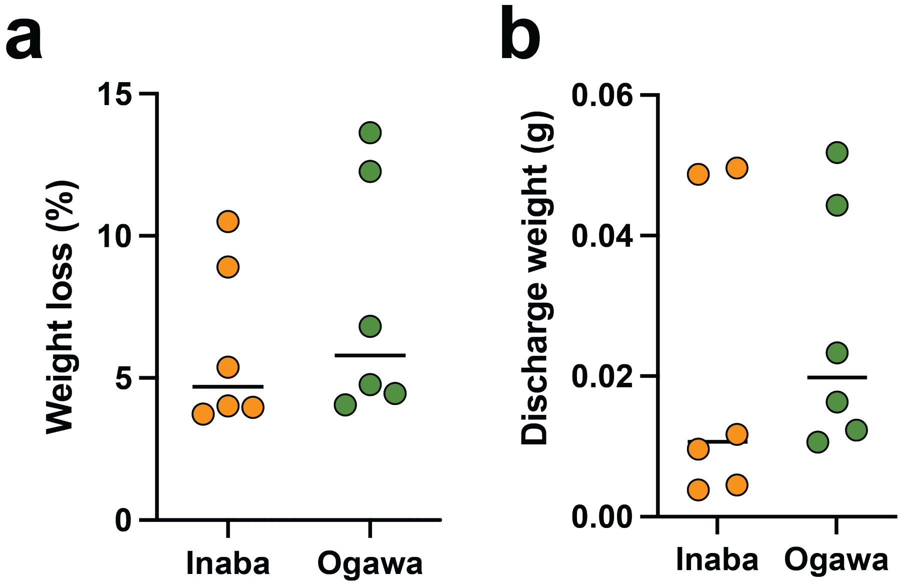


Supplementary Figure 2: **Measurements of weight loss and diarrheal discharge.** Weight loss (c) and weight of diarrheal discharge (d) of p5 suckling mice after 24 of infection with Inaba (orange) or Ogawa (green) serotype. All mice used were Crl:CD1(ICR) mixed sex, postnatal day 5 at the time of infection (n = 6). Source data are provided as a Source Data file.

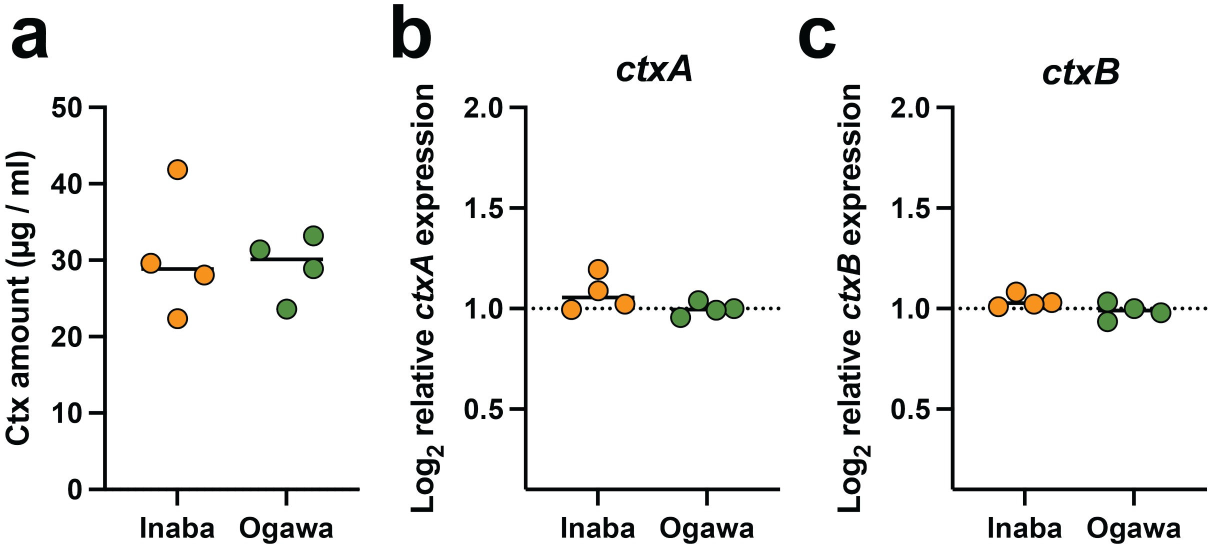


Supplementary Figure 3: **Impact of serotype on cholera toxin expression**. (a) amount of cholera toxin was measured by CTX ELISA of the Inaba (orange) or Ogawa (green) serotype grown in AKI medium (n = 4). qRT-PCR measurements of ctxA (b) and ctxB (c) of the Inaba (orange) or Ogawa (green) serotype after 18 h of infection of suckling mice (n = 4). All mice used were Crl:CD1(ICR) mixed sex, postnatal day 5 at the time of infection. Source data are provided as a Source Data file.

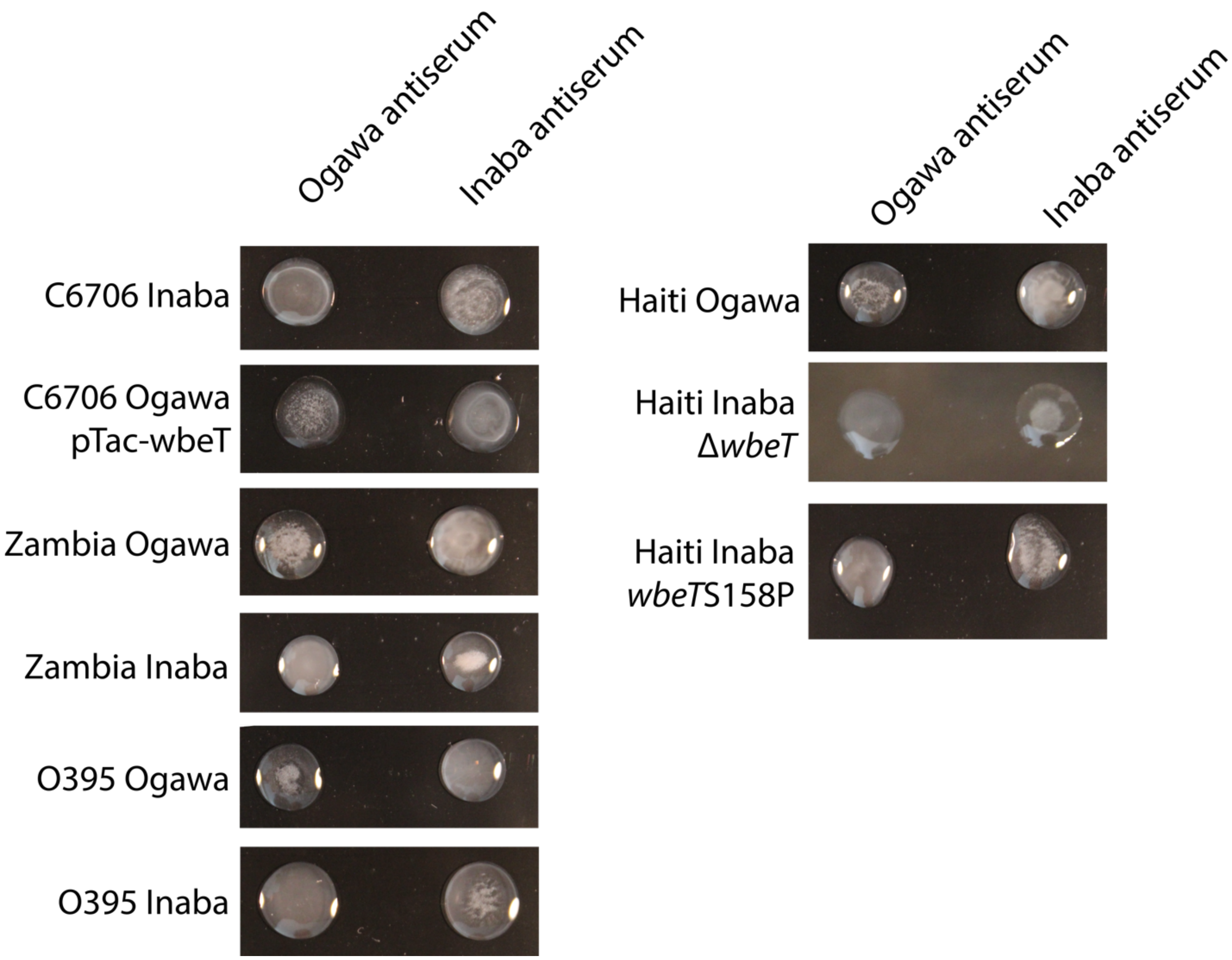


Supplementary Figure 4: **Serotyping of *V. cholerae* isolates used in this study**. *V. cholerae* cultures treated with Ogawa or Inaba specific antisera. Images were taken after 5-10 minutes of incubation at room temperature.

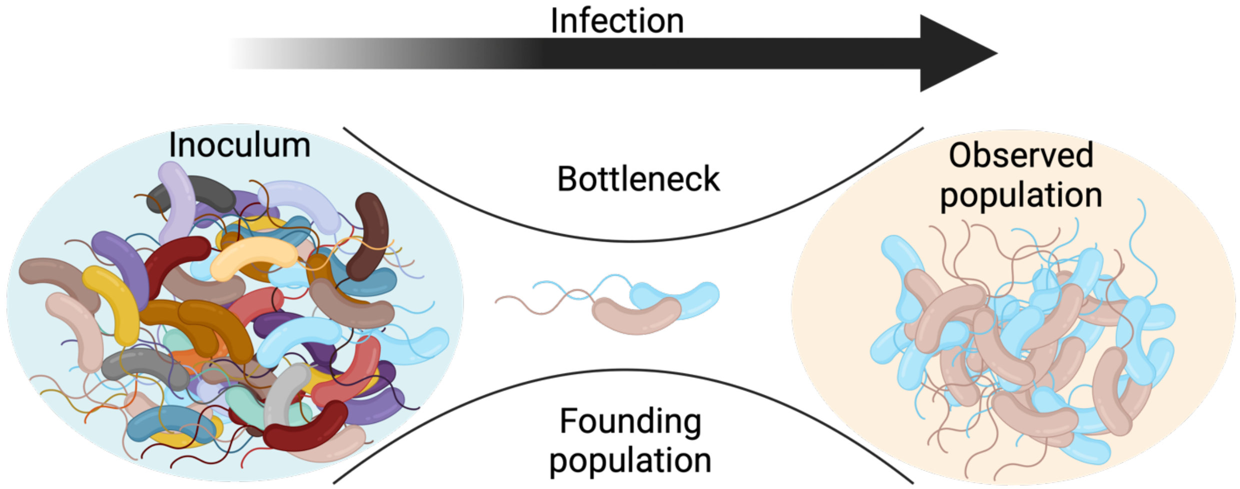


Supplementary Figure 5: **Schematic depicting assays to measure founding populations.** Barcoded bacteria are used to measure the number of bacteria that survive the bottleneck, the founders, and give rise to the population in intestinal homogenates. Parts of this figure were created in BioRender. Zingl, F. (2025) https:// BioRender.com/fhbbff0.

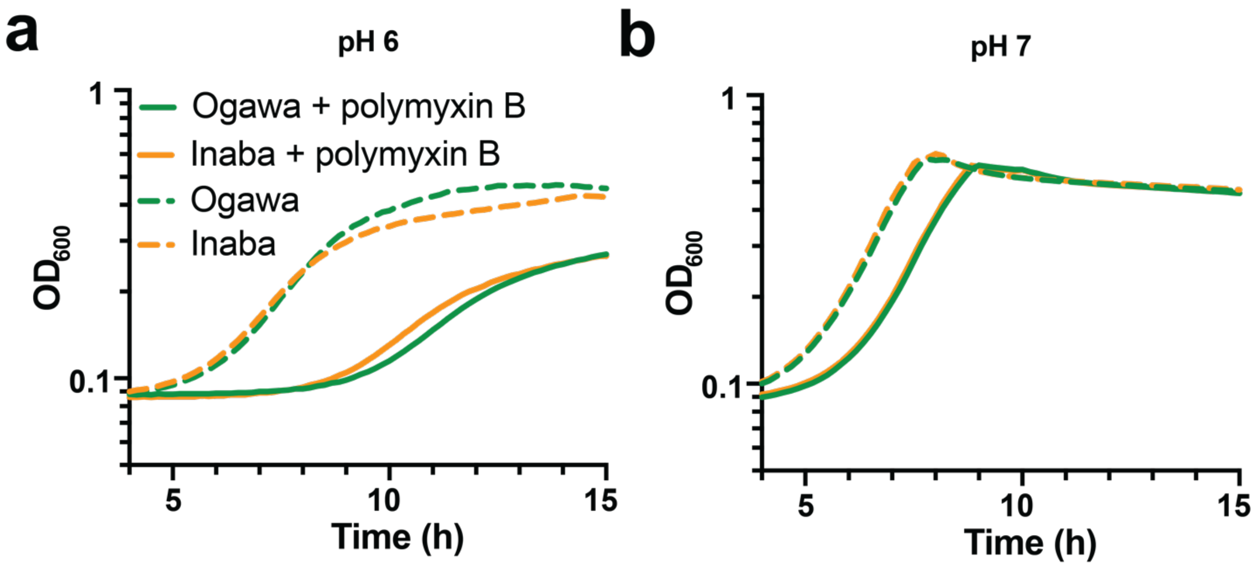


Supplementary Figure 6: **Growth curves at pH 6 and pH 7.** Growth curves at pH 6 (a; n = 3) and pH 7 (b; n = 4) of Inaba (orange) or Ogawa (green) strains in M9 with (solid line) or without (dashed line) polymyxin B. Source data are provided as a Source Data file.
